# H3K4me3 instructs transcription at intergenic active regulatory elements

**DOI:** 10.1101/2025.03.21.644682

**Authors:** Haoming Yu, Yongyan Zhang, Zhicong Liao, Benjamin William Walters, Bluma J. Lesch

## Abstract

Mammalian genomes undergo pervasive transcription in both genic and intergenic regions. Tri-methylation of histone H3 lysine 4 (H3K4me3) is a deeply conserved and functionally important histone modification enriched at transcriptionally active promoters, where it facilitates RNA polymerase activity. H3K4me3 is also commonly found in intergenic regions, where its role is poorly understood. We interrogated the role of H3K4me3 at intergenic regulatory elements by using epigenetic editing to efficiently deposit H3K4me3 at intergenic loci. We find that H3K4me3 instructs RNA polymerase activity and is actively remodeled at intergenic regions, shedding light on these important but poorly understood regions of the genome.

## Introduction

Methylation of lysine 4 on histone H3 (H3K4me) is a widespread, evolutionarily conserved, and extensively studied histone modification. H3K4 can gain one, two, or three methyl groups, and each of the three H3K4 methylation states (mono-, di-, or trimethylation, H3K4me1/2/3) is associated with different genomic features. Specifically, H3K4me1 is a marker for enhancer regions, while H3K4me3 is a marker for transcriptionally active gene promoters. H3K4 methyltransferase and demethylase protein complexes are essential for cell differentiation and development and are frequently mutated in cancers, implying that regulation of H3K4 methylation is important for biological function.^1-3^ Due to their importance in cellular function and disease, H3K4me1/2/3 modifications are among the most studied and best understood histone modifications. However, there remain major unsolved problems regarding their functional roles and the reasons for their specific and distinct genomic distributions.

The level of enrichment of H3K4me3 is highly correlated with level of transcription initiation and pause release, and H3K4me3 interacts directly with transcriptional cofactor TAF3, as well as chromatin remodelers such as BPTF and CHD1.^4-7^ Knockouts and catalytic domain mutations of H3K4 methyltransferases and demethylases suggest that H3K4me3 is involved in transcriptional initiation and elongation in a context-dependent manner, and targeted rapid degradation of H3K4 tri-methyltransferase complexes supported an active role for the H3K4me3 modification in promoter-proximal pause release.^8-13^ However, noncatalytic functions of the enzymes, indirect effects from genome-wide transcriptional changes and potential compensation effects from other H3K4 tri-methyltransferase complexes mean that other mechanisms contributing to transcription beyond H3K4me3-dependent signaling could not be fully excluded. To avoid these indirect effects, the functional role of H3K4me3 at specific promoter loci has been interrogated using epigenome editing with a targeted catalytically dead Cas9 (dCas9) fused to a H3K4 methyltransferase domain. Deposition of H3K4me3 at candidate promoters using this approach was sufficient to increase transcript levels when DNA methylation levels were low, but could not overcome a DNA hypermethylated state.^14,15^ These findings at promoters highlight a significant open question: what directs H3K4me3-mediated transcriptional activity in intergenic regions of the genome? H3K4me3 is enriched at a subset of transcriptionally active enhancers^16-19^, suggesting that it may have a general role in transcriptional initiation throughout the genome with a broad potential to regulate hundreds of thousands of active, primed, and dormant cis-regulatory regions. A deeper understanding of H3K4me3 and transcriptional regulation at these important regions of the genome is crucial for understanding the mechanisms of H3K4 modifying enzymes and the drugs that target their activities in cancer and other disease states.

Here, we hypothesized that H3K4me3 is sufficient to support transcription even at distal regulatory elements. We generated and applied an epigenetic editing tool, dCas9-PRDM9, systematically installed H3K4me3 at varied intergenic candidate cis-regulatory elements (cCREs) in human cells, and examined the consequences for local transcription and enhancer function in order to define genome-wide, context-specific functions for H3K4me3.^14,15^ We found that H3K4me3 is sufficient to increase transcription at active cCREs, regardless of distance to the nearest gene or physical interaction with a promoter region. This effect was independent of cCRE identity or location, unlinked from transcript levels of putative target genes, and dependent on chromatin context. Our results demonstrate a generalizable instructive role for H3K4me3 in supporting transcription throughout the genome.

## Results and Discussion

To understand how H3K4me3 is controlled at putative enhancers compared to promoters, we first examined the impact of disrupting H3K4 tri-methyltransferase or H3K4 demethylase activity on H3K4me3 distributions at intergenic regions and promoters. We reanalyzed published H3K4me3 ChIP-seq data from cells lacking *RACK7*, a cofactor for the KDM5C H3K4 demethylase, or for *Cfp1*, a subunit of the SET1 H3K4 tri-methyltransferase complex.^20-22^ In wild type cells, H3K4me3 was already present at many intergenic regions. However, disrupting either demethylase or methyltransferase functions caused significant new H3K4me3 deposition at intergenic regions, and this effect impacted intergenic sites substantially more than gene promoters. This result suggests that H3K4me3 is continuously removed from intergenic sites by the RACK7/KDM5C demethylase machinery and also continuously guided to promoters by CFP1/SET1 (**Figure 1A, Supplemental Table 1, Supplemental Table 2**). Thus, accumulation of H3K4me3 at intergenic sites is widespread but appears to be actively remodeled under normal conditions.

**Figure 1.**
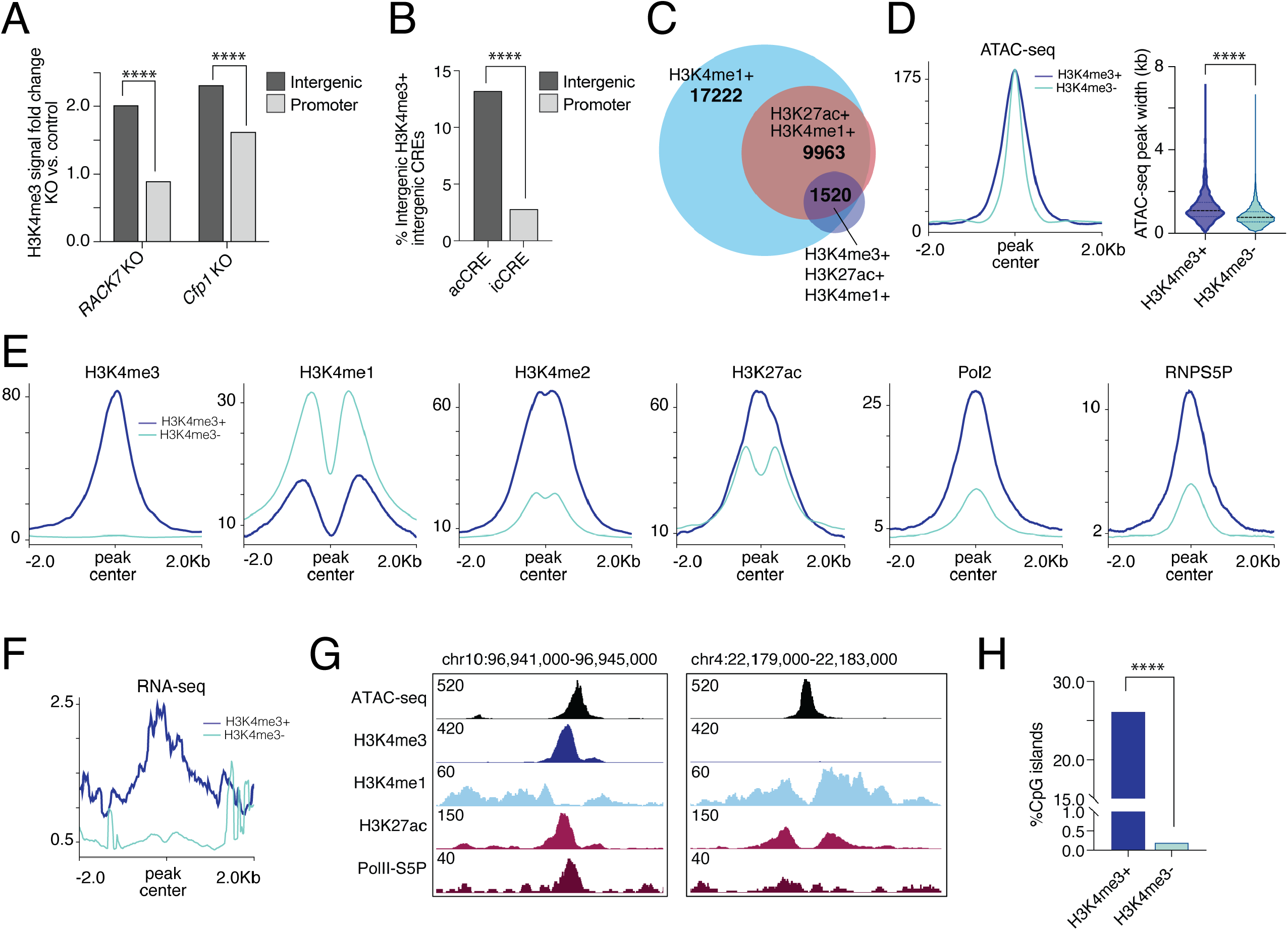
Intergenic cCREs enriched for H3K4me3 are associated with higher transcriptional activity. **A**, Fold change (knockout vs. wild type) of numbers of H3K4me3 ChIP-seq peaks at intergenic and promoter regions in *RACK7* knockout human breast cancer (MDA-MB-231) cells or *Cfp1* knockout mouse embryonic stem cells. \*\*\*\**P* < 0.0001, Fisher’s exact test. **B**, Percentage of intergenic acCREs and icCREs that overlap H3K4me3 ChIP-seq peaks. \*\*\*\**P* < 0.0001, Fisher’s exact test. **C**, Sets of intergenic ATAC-seq peaks (total n=44,832) that overlap with H3K4me1 only (n=17,222), H3K27ac and H3K4me1 (n=9,963), or H3K4me3, H3K27ac, and H3K4me1 (n=1,520) ChIP-seq peaks in MDA-MB-231 cells. The latter two groups (n=11,483 total) are considered acCREs. **D**, Metagene plot of ATAC-seq signal (left) and comparison of ATAC-seq peak widths (right) for H3K4me3+ and H3K4me3-acCREs. \*\*\*\**P* < 0.0001, Welch’s t test. **E**, Metagene plots of H3K4me3, H3K4me1, H3K4me2, H3K27ac, total Pol II, and Pol II-S5P ChIP-seq reads at H3K4me3+ and H3K4me3-acCREs. **F**, Metagene plot of RNA-seq reads at H3K4me3+ and H3K4me3-acCREs. **G**, Genome browser tracks for ChIP-seq signal at representative H3K4me3+ and H3K4me3-acCREs. **H**, Percentage of acCREs and icCREs that overlap CpG islands. \*\*\*\**P* < 0.0001, Fisher’s exact test.

We next set out to determine the functional implications of intergenic H3K4me3 deposition. In order to precisely map intergenic H3K4me3 peaks and define their distinct chromatin features, we performed H3K4me3 ChIP-seq in human breast cancer cells, and classified H3K4me3 peaks as intergenic based on a distance of at least 2 kilobases from annotated genes. Comparison to RNA-seq data from the same cell type confirmed that these intergenic peaks did not overlap unannotated transcripts. To characterize the chromatin context of intergenic H3K4me3 peaks, we collected genome-wide profiles of H3K4me1, H3K4me2, H3K27ac, total Pol II, and Pol II-S5P (Pol II phosphorylated on Serine 5, a feature of initiating Pol II) using ChIP-seq, as well as ATAC-seq and RNA-seq data from the same cells (**Supplemental Figure 1**). Consistent with previous reports, we found that intergenic H3K4me3 peaks were significantly enriched at active candidate cis-regulatory elements (acCREs, defined as intergenic ATAC-seq peaks that overlap both H3K27ac and H3K4me1 peaks), compared to inactive candidate cis-regulatory elements (icCREs, defined as intergenic ATAC-seq peaks lacking H3K27ac or H3K4me1) **(Figure 1B, Supplemental Table 3**).

To determine if the presence of H3K4me3 is associated with a functionally distinct subset of acCREs, we next subdivided acCREs into H3K4me3+ (n=1520), and H3K4me3-(n=9963) for further analysis **(Figure 1C**). Overall, H3K4me3+ acCREs were slightly broader than H3K4me3-acCREs with similar peak chromatin accessibility (**Figure 1D**). Enrichment of H3K4me3 at acCREs was associated with lower H3K4me1, higher H3K4me2, higher H3K27ac, stronger total Pol II binding, higher levels of initiating Pol II (Pol II-S5P), and higher transcript levels (**Figure 1E-F**). Interestingly, H3K27ac signal tended to be unimodal and correlated with the central H3K4me3 peak at H3K4me3+ acCREs, but was bimodal and more closely correlated with H3K4me1 signal at H3K4me3-acCREs (**Figure 1E, 1G**). About one fourth of H3K4me3+ acCREs included CpG islands, while almost no H3K4me3-acCREs included annotated CpG islands, suggesting that recruitment of SET1/MLL H3K4 methyltransferase complexes to intergenic CpG islands may bias these sites toward accumulation of H3K4me3 by helping to overcome active H3K4 demethylation (**Figure 1H**). However, the presence of CpG islands does not explain the presence of H3K4me3 at the majority of H3K4me3+ acCREs. Overall, we found that only a small portion of intergenic acCREs is enriched with H3K4me3 but that this subset has distinct genomic and epigenomic features. Specifically, H3K4me3+ acCREs have features associated with higher transcriptional activity, consistent with a well-established relationship between H3K4me3 deposition and transcription.

H3K4me3 enrichment is correlated with features of active transcription at acCREs. We hypothesized that H3K4me3 plays a functional role in boosting transcription at these sites. To directly test the functional role of H3K4me3 in a locus-specific manner, we generated an epigenetic editing tool, dCas9-PRDM9, by fusing the catalytic (SET) domain of the H3K4 tri-methyltransferase PRDM9 to a catalytically dead Cas9 (dCas9) protein. The PRDM9 catalytic domain has been shown to be sufficient for mono-, di-, and tri-methylation of unmodified H3K4, independent of additional cofactors, and to mediate locus-specific H3K4me3 deposition in a multiplexed epigenetic editing system.^14,15,23^ As a control, we generated an identical fusion protein with a single point mutation abrogating methyltransferase catalytic activity (dCas9-cdPRDM9) (**Figure 2A**). Unlike knockout studies where deletion of histone modifiers removes both catalytic and non-catalytic functions, this tool allowed us to deposit H3K4me3 at a specific acCRE and study transcriptional changes both at the acCRE itself and its interacting gene promoter while controlling all other aspects of the system. Total H3K4me1 and H3K4me3 levels, as well as expression levels of the fusion proteins, were comparable in stable cell lines expressing dCas9-PRDM9 or dCas9-cdPRDM9 (**Supplemental Figure 2A**).

**Figure 2.**
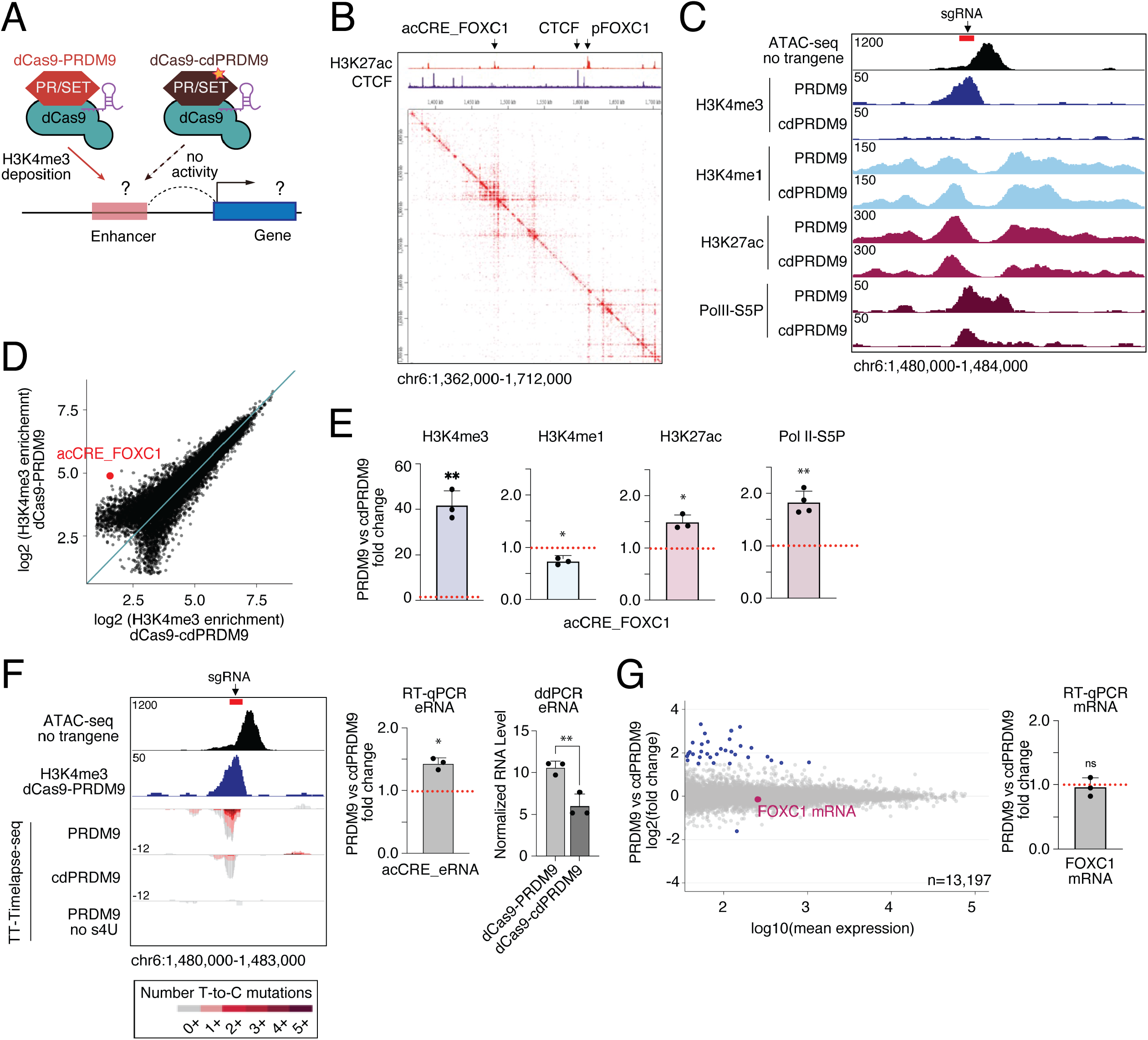
H3K4me3 facilitates local transcription at an isolated acCRE. **A**, Schematic of dCas9-PRDM9 and dCas9-cdPRDM9 and experimental design for studying functional roles of H3K4me3 at enhancers. **B**, Genome browser tracks of H3K27ac and CTCF ChIP-seq signals (top), and heatmap representing long-range chromatin contact probabilities measured by H3K27ac HiChIP in wild type MDA-MB-231 cells in the region of acCRE_FOXC1. Arrows indicate acCRE_FOXC1, *FOXC1* gene promoter, and insulating CTCF binding site. Data from GEO:GSM2572593. **C**, Genome browser tracks for ATAC-seq in wild type cells, and normalized H3K4me3, H3K4me1, H3K27ac, and Pol II-S5P ChIP-seq signals in dCas9-PRDM9 and dCas9-cdPRDM9 cells at acCRE_FOXC1. Arrow indicates sgRNA site. **D**, Scatter plot of normalized H3K4me3 ChIP-seq reads at all H3K4me3 peaks in dCas9-PRDM9 and dCas9-cdPRDM9 cells. Red point represents the deposited H3K4me3 peak at acCRE_FOXC1 (called as a peak only in the dCas9-PRDM9 condition). **E**, Fold change of H3K4me3, H3K4me1, H3K27ac, and Pol II-S5P signal in dCas9-PRDM9 compared to dCas9-cdPRDM9 cells at the acCRE_FOXC1 locus measured by ChIP-qPCR. \**P* < 0.05, \*\**P* < 0.01, one-sample t test. **F**, Left, genome browser tracks for ATAC-seq in wild type cells, normalized H3K4me3 ChIP-seq signal, and nascent transcript reads as measured by TT-Timelapse-seq in dCas9-PRDM9 and dCas9-cdPRDM9 cells at the acCRE-FOXC1 locus. Red bar indicates sgRNA site. Color of the nascent transcript signal stands indicates the number of T-to-C mutations found per read by TT-Timelapse-seq. Fold changes of acCRE_FOXC1 RNA levels measured by RT-qPCR (center) and RT-droplet digital PCR (RT-ddPCR) (right) are shown. \**P* < 0.05, \*\**P* < 0.01, unpaired t test for RT-ddPCR, one-sample t test for RT-qPCR. **G**, Left, MA plot of total transcriptome (RNA-seq) reads in dCas9-PRDM9 compared to dCas9-cdPRDM9 (n=2 biological replicates). Right, fold change of FOXC1 mRNA by RT-qPCR. ns, not significant at a threshold of *P* < 0.05, one-sample t test. Error bars represent mean ±SD (n=3-4 biological replicates).

To validate the efficiency of dCas9-PRDM9, we selected an isolated H3K4me3-negative acCRE located 130kb upstream of its nearest active gene *FOXC1* (acCRE_FOXC1). While acCRE_FOXC1 was previously proposed as a super-enhancer for *FOXC1*, it was later found to be insulated from physical interaction with the *FOXC1* promoter by CTCF (**Figure 2B**).^24,25^ This acCRE is therefore an optimal site to study the local effects of H3K4me3 without feedback from promoter interactions and coding transcripts. We stably co-expressed dCas9-PRDM9 and an sgRNA targeting the ATAC-seq signal peak of acCRE_FOXC1, and quantified changes in chromatin state by ChIP-qPCR and sequencing (**Supplemental Figure 2B-C**). H3K4me3 was efficiently deposited at the target locus with minimal off-target effects and at levels comparable to H3K4me3+ acCREs (**Figure 2C-D, Supplemental Figure 2D-F, Supplemental Table 4**). H3K4me3 deposition was efficient enough to reduce H3K4me1 levels at the target locus, as the PRDM9 catalytic domain drives conversion of H3K4me1 to H3K4me3 (**Figure 2C, 2E**). Deposition of H3K4me3 also led to modest gains in H3K27ac (**Figure 2C, 2E**). Meanwhile, there was a substantial increase in initiating Pol II-S5P (**Figure 2C, 2E**), suggesting increased transcriptional activity. Indeed, local transcription was also consistently upregulated as assayed by TT-Timelapse-seq, RT-ddPCR, and RT-qPCR (**Figure 2F, Supplemental Figure 2G**). Nascent transcription assays also localized the gain in H3K4me3 to the apparent +1 nucleosome relative to transcriptional initiation, and suggest that the initiated transcript is terminated relatively quickly, reflecting features of an enhancer RNA (eRNA) (**Figure 2F**). In contrast, no transcriptomic changes were seen at the global level, and levels of *FOXC1* mRNA were unchanged (**Figure 2G, Supplemental Figure 2H, Supplemental Table 5**). Together, these results indicate that H3K4me3 is sufficient to facilitate local transcription of eRNAs at an isolated acCRE, independent of interactions with an established transcriptional start site.

Next, we asked if deposition of H3K4me3 at acCREs could impact enhancer activity. We directed H3K4me3 to two acCREs previously shown to interact with active gene promoters: a proximal enhancer located 14kb upstream of gene CSF1 (eCSF1), and a distal enhancer located 180kb upstream of gene FOXQ1 (e2_FOXQ1). We generated individual cell lines carrying the dCas9-PRDM9 or dCas9-cdPRDM9 constructs and a sgRNA targeting one of these two enhancers. H3K4me3 was efficiently and specifically installed at the apparent +1 nucleosome at both enhancers (**Figure 3A**). Consistent with our results at acCRE_FOXC1, H3K27ac and initiating Pol II-S5P were significantly increased following deposition of H3K4me3 at both loci (**Figure 3B**). eRNA transcript levels also increased at both loci (**Figure 3C**). However, we did not detect any change in mRNA transcript levels of the target coding genes, suggesting that H3K4me3 and consequent transcription at enhancers did not alter enhancer activity (**Figure 3D**). Overall, we found that H3K4me3 is sufficient to facilitate local transcription at active enhancers, but is dispensable for enhancer activity.

**Figure 3.**
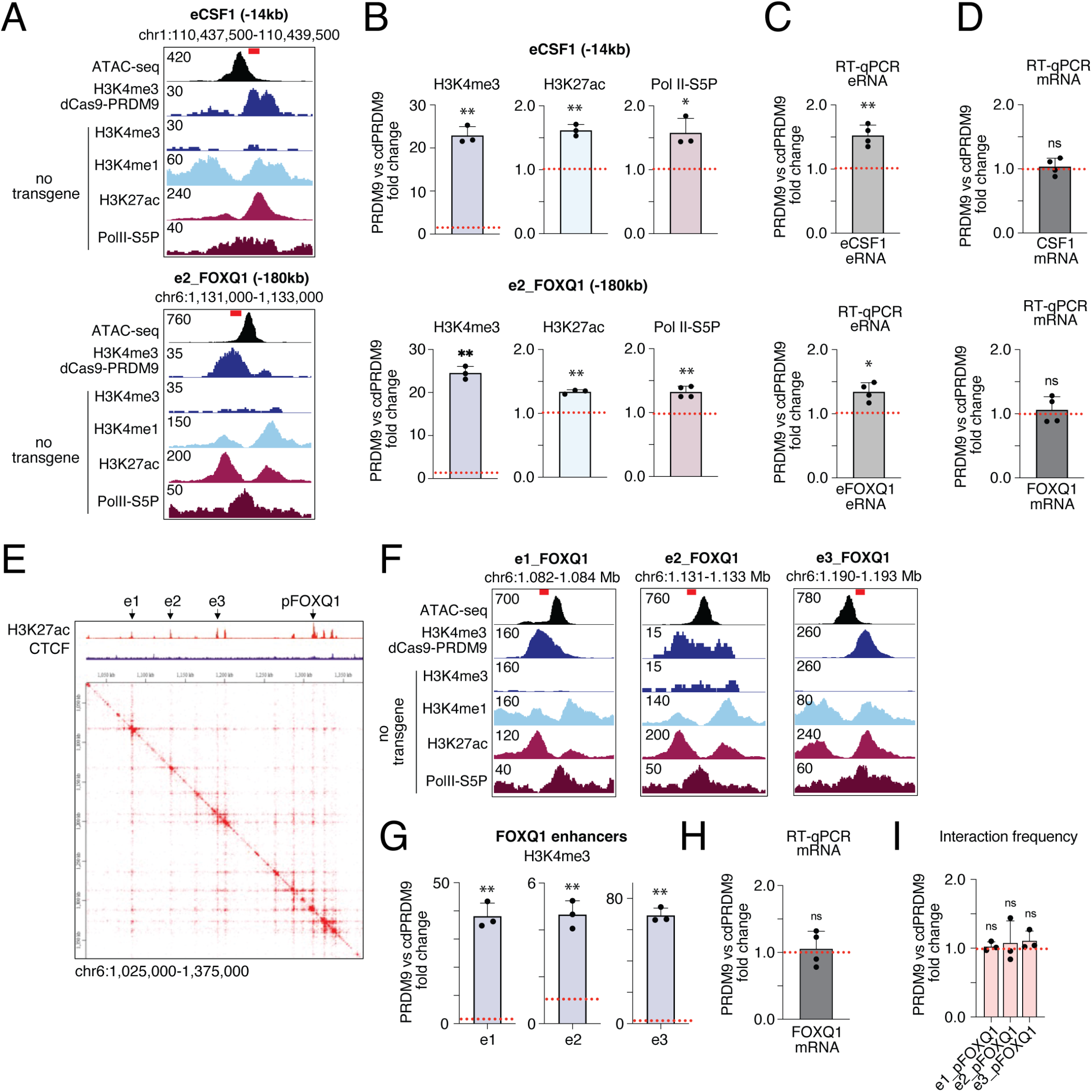
H3K4me3 boosts eRNA transcription at active enhancers but is dispensable for enhancer activity. **A**, Genome browser tracks for ATAC-seq, H3K4me3, H3K4me1, H3K27ac, and Pol II-S5P ChIP-seq signals in wild type cells, and H3K4me3 ChIP-seq signals in dCas9-PRDM9 cells at the eCSF1 and e2_FOXQ1 loci. Red bars indicate sgRNA sites. **B**, Fold change for H3K4me3, H3K27ac, and Pol II-S5P levels in dCas9-PRDM9 compared to dCas9-cdPRDM9 cells at eCSF1 and e2_FOXQ1 by ChIP-qPCR. \**P* < 0.05, \*\**P* < 0.01, one-sample t test. **C**, Fold change for eCSF1 and e2_FOXQ1 eRNA levels measured by RT-qPCR. \**P* < 0.05, \*\**P* < 0.01, one-sample t test. **D**, Fold change for *CSF1* and *FOXQ1* mRNA levels measured by RT-qPCR. ns, not significant at a threshold of *P* < 0.05, one-sample t test. **E**, Genome browser tracks for H3K27ac and CTCF ChIP-seq signals (top), and heatmap representing long-range chromatin contact probabilities measured by H3K27ac HiChIP in wild type MDA-MB-231 cells in the region of e2_FOXQ1. Arrows point to three target *FOXQ1* enhancers and the *FOXQ1* gene promoter. Data from GEO:GSM2572593. **F**, Genome browser tracks for ATAC-seq, H3K4me3, H3K4me1, H3K27ac, and Pol II-S5P ChIP-seq signals in wild type cells, and H3K4me3 ChIP-seq signal in dCas9-PRDM9 cells at three target FOXQ1 enhancers. e2_FOXQ1 corresponds to the enhancer shown in A-D. Red bars indicate sgRNA sites. **G**, Fold change of H3K4me3 at three target *FOXQ1* enhancers in dCas9-PRDM9 compared to dCas9-cdPRDM9 measured by ChIP-qPCR. ** *P* < 0.01, one-sample t test. **H**, Fold change of *FOXQ1* mRNA levels measured by RT-qPCR. ns, not significant at a threshold of *P* < 0.05, one-sample t test. **I**, Fold change of normalized interaction frequencies between target enhancers and the *FOXQ1* promoter measured by 3C-qPCR. ns, not significant at a threshold of *P* < 0.05, one-sample t test. Error bars represent mean ±SD (n = 3-4 biological replicates).

In eukaryotes, multiple enhancers can form a transcription initiation hub to regulate one gene promoter. Since deposition of H3K4me3 at one enhancer did not change its target gene transcription, we wondered whether deposition of H3K4me3 at multiple interacting enhancers simultaneously could upregulate target gene transcription. We generated cell lines carrying the dCas9-PRDM9 or dCas9-cdPRDM9 constructs and a multiplex sgRNA expression cassette targeting multiple enhancers shown to interact with the promoters of FOXQ1 or another active gene, SOX9 (**Figure 3E, Supplemental Figure 3**). H3K4me3 was specifically installed at the apparent +1 nucleosome for all target enhancers with either single or multiplexed sgRNAs, although efficiency of H3K4me3 deposition was variable between different enhancer elements (**Figure 3F-G, Supplemental Figure 3A,C-D**). Deposition of H3K4me3 at multiple enhancers simultaneously did not alter target gene mRNA levels or enhancer-promoter interaction frequencies at each locus, again supporting a model where transcription at enhancers does not directly impact transcription of the target promoter (**Figure 3H-I, Supplemental Figure 3E-F**).

Our results indicate that H3K4me3 is sufficient to instruct transcription at acCREs. acCREs are already in a transcriptionally accessible chromatin state, raising the question of whether H3K4me3 is also sufficient to initiate transcription at icCREs. To address this, we divided icCREs into two groups: primed enhancers (PE, defined as intergenic ATAC-seq peaks that overlap H3K4me1 peaks but not H3K27ac peaks), and unmodified open chromatin (UO, defined as intergenic ATAC-seq peaks that do not overlap either H3K4me1 or H3K27ac peaks). We used dCas9-PRDM9 to direct H3K4me3 to two primed enhancers (PE1, PE2) and two unmodified open chromatin regions (UO1, UO2) in separate cell lines. Similar to acCREs, H3K4me3 was efficiently and specifically deposited at the nearest nucleosome at each target locus (**Figure 4A-B, Supplemental Figure 4A-B**). However, deposition of H3K4me3 did not promote substantial gain of either H3K27ac or initiating Pol II-S5P at either PEs or UOs, although there was a mild increase in H3K27ac at both PEs (**Figure 4C, Supplemental Figure 4C**). The differential effects of H3K4me3 deposition at acCREs compared to icCREs indicate that H3K4me3 is not sufficient to instruct transcriptional activation alone. Instead, H3K4me3 facilitates transcription in the context of prior transcriptional machinery and acetylated nucleosomes (**Figure 4D**). Thus, H3K4me3 is sufficient to boost local transcription at acCREs, and this activity is dependent on chromatin context but independent of target gene activity.

**Figure 4.**
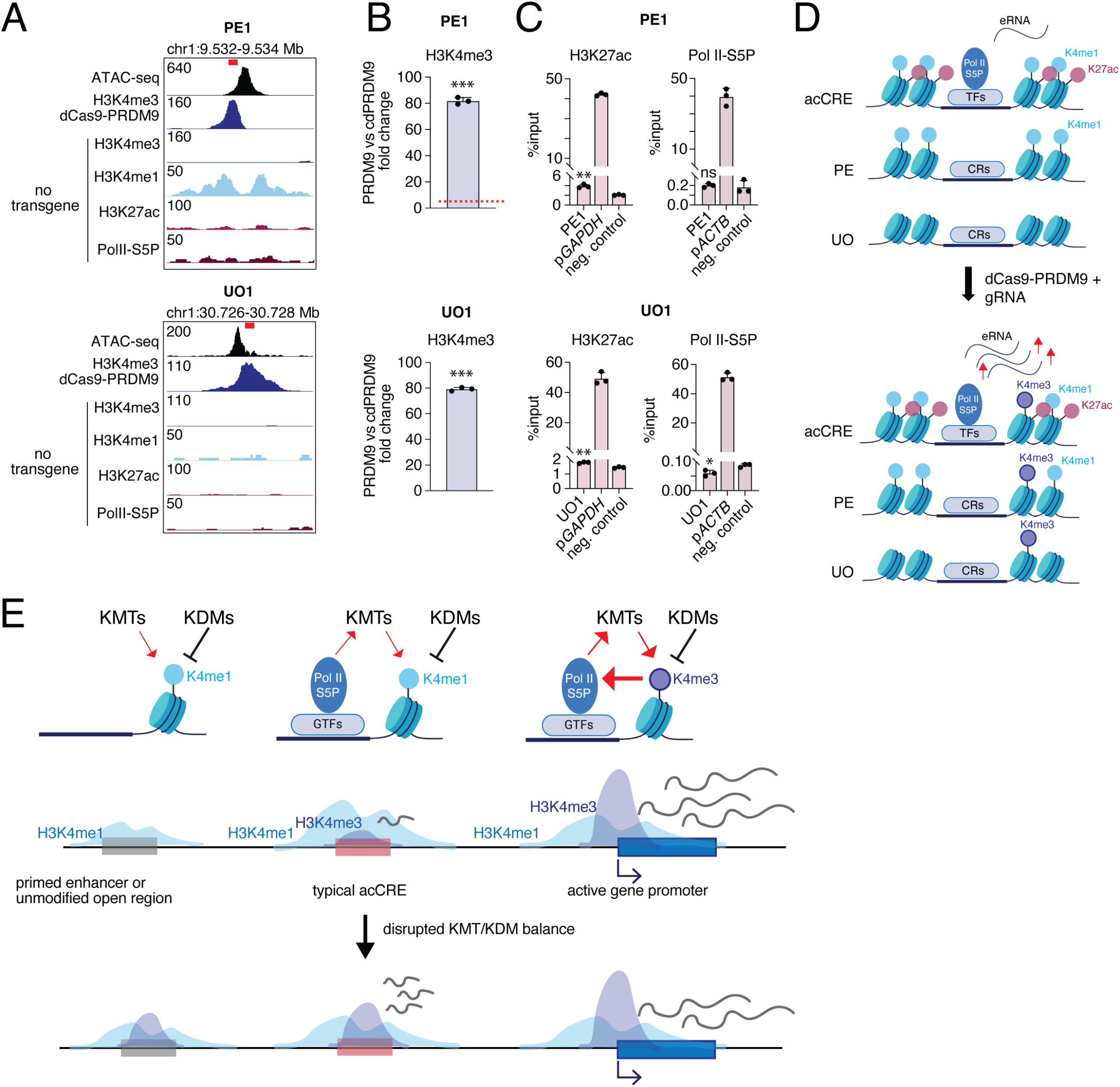
Instruction of transcription by H3K4me3 at cCREs depends on chromatin context. **A**, Genome browser tracks for ATAC-seq, H3K4me3, H3K4me1, H3K27ac, and Pol II-S5P ChIP-seq signals in wild type cells, and H3K4me3 ChIP-seq signals in dCas9-PRDM9 cells at the PE1 and UO1 loci. Red bars indicate sgRNA sites. **B**, Fold change in H3K4me3 at PE1 and UO1 loci in dCas9-PRDM9 compared to dCas9-cdPRDM9 cells measured by ChIP-qPCR. \*\*\**P* < 0.001, one-sample t test. **C**, Relative enrichment of H3K27ac and Pol II-S5P at a positive control region (*GAPDH* or *ACTB* gene promoter), a negative control region (gene desert locus on chromosome 12), PE1, and UO1 measured by ChIP-qPCR. \**P <* 0.05, \*\**P <* 0.01, ns, not significant at a threshold of *P* < 0.05, unpaired t-test compared to negative control. Error bars in B and C represent mean ±SD (n = 3 biological replicates). **D**, Schematic diagram showing differential outcomes of H3K4me3 deposition at acCREs and icCREs (PE and OU). **E**, Proposed model showing remodeling of H3K4me3 away from intergenic enhancers and toward promoter regions under normal conditions, and changes in distribution when methyltransferase (KMT) /demethylase (KDM) balance is disrupted. Transcriptional effects at promoters are variable in the disrupted context.

Together, our results indicate that H3K4me3 is not required for enhancer activity, but is sufficient to promote transcription in a permissive chromatin environment. We suggest that H3K4me3 is actively depleted from intergenic cCREs in favor of H3K4me1 in order to more efficiently direct transcriptional activity to licensed gene promoters and prevent noisy transcription at alternative sites. More broadly, this suggests that regulation of intergenic transcription plays an important guardian function to promote robust and stable transcriptional states across the genome. Adaptation of the dCas9-PRDM9 tool to a modular, scalable system would allow us to test this model by targeting many loci simultaneously across multiple cell types.

Our results also raise several unresolved questions. First, while we did not observe a gain in target gene mRNA transcription when we drove increased eRNA transcription from acCREs with H3K4me3, previous reports have shown that strongly activating epigenome editors such as dCas9-p300 and dCas9-VP64 (CRISPRa) can increase both eRNA and target gene mRNA levels when recruited to an enhancer locus.^26-29^ We see two principal possible explanations for this difference: first, recruitment of a much stronger transcriptional activator such as p300 or VP64 could increase eRNA level above a certain threshold not obtained by H3K4me3 in our study, and this level of eRNA transcription may be sufficient to enhance transcription at a target promoter. Second, p300 or VP64 might increase the enhancer activity of the target cCRE via a mechanism independent of the effects on eRNA transcription. For example, acetylated nucleosomes could recruit general transcription factors (GTFs) to form a more concentrated transcription initiation hub to mediate target gene transcription activation through promoter-enhancer contacts.^30,31^ An additional possibility is that compensation by other, unidentified enhancer regions maintains stable levels of the target gene transcript, although we did not see evidence of this effect at the FOXQ1 locus. Future work will be important to distinguish between these possibilities.

Second, our study does not fully resolve the mechanism by which H3K4me3 boosts transcription at acCREs. Based on previous interaction studies, a likely mechanism is that H3K4me3 enhances transcription through interaction with TAF3, a subunit of TFIID, to stabilize the transcription initiation complex.^6,7^ A role in stabilizing initiation at enhancers could represent a qualitative difference from H3K4me3 function at promoters, where recent work suggests it may function in Pol II pause release.^12^ Although we were unable to directly test this mechanism due to lack of a ChIP-grade TAF3 antibody, we did observe an increase of TBP binding at target sites, as predicted if H3K4me3 recruits TAF3. Direct evaluation of this model will be important to fully understand how H3K4me3 impacts transcription and if this function differs at enhancer compared to promoter regions.

Finally, if H3K4me3 is actively depleted from intergenic cCREs, why is it still present at a subset of these regions? The observed level of H3K4me3 at a given locus represents the output of a balance between H3K4 methyltransferases and demethylases. H3K4me3 level is itself partly dependent on transcription, as blocking transcription initiation leads to a dramatic loss of H3K4me3 at promoters and enhancers.^32^ Thus, H3K4 tri-methyltransferase binding may be stabilized by active transcription, promoting H3K4me3 deposition and in turn facilitating transcription as a feedback loop. This feedback likely allows H3K4me3 methyltransferase activity to outbalance demethylase activity at active promoters, and may also allow H3K4me3 to accumulate at the most highly transcribed acCREs, escaping the general mechanism directing H3K4me3 to promoters. In contrast, at the majority of enhancer regions, lower-level RNA polymerase activity and early termination of transcription tip the balance toward demethylase activity, ensuring H3K4me3 depletion at enhancers to tune down surplus transcriptional activity (**Figure 4E**). Active depletion of H3K4me3 at intergenic cCREs may help to guide strong transcriptional activity to promoter regions or avoid unwanted initiation of cryptic transcripts, promoting a robust and stable transcriptional state.

## Material and Methods

### Cell culture

MDA-MB-231 cells (ATCC, HTB-26) and 293T cells (ATCC, CRL-3216) were grown in complete DMEM media (Gibco) supplemented with 10% fetal bovine serum (Sigma‐Aldrich), 1:100 Penicillin-Streptomycin (Gibco), and 2.5 ug/ml Plasmocin (InvivoGen). Cells were grown at 37°C at 5% CO2, and passaged when they reached 70–80% confluence. Cells were screened monthly for mycoplasma and results were excluded if cells were found to be mycoplasma positive.

### Antibodies

The following antibodies were used: H3K4me1 (Cell Signaling Technology, 5326), H3K4me2 (Abcam, ab7766), H3K4me3 (EpiCypher, 13-0041), H3K27ac (Abcam, ab4729), Total Pol-II (Diagenode, C15200004), and Pol II-S5P (Cell Signaling Technology, 13523) for ChIP. H3K4me1 (Abcam, ab8895), H3K4me3 (Abcam, ab8580), pan-H3 (Abcam, ab1791), GAPDH (Santa Cruz Biotechnology, sc32233), CRISPR/Cas9 (Diagenode, C15310258), goat anti‐rabbit IgG conjugated to HRP (Jackson Immuno Research), and goat anti‐mouse IgG conjugated to HRP (Jackson Immuno Research) for Western blotting.

### Fusion protein and sgRNA cloning

The mouse PRDM9 cDNA was a kind gift from Dr. Galina Petukhova. The PRDM9 catalytic domain cDNA (position 574-1332 of the cDNA) was cloned into a dCas9 fusion vector (Addgene, 51023). The Q5 Site-Directed Mutagenesis Kit (NEB) was used to introduce the catalytic point mutation to generate dCas9-cdPRDM9. dCas9-PRDM9/cdPRDM9 was first cloned into the pSBbi-Neo plasmid (Addgene, 60525) to add a neomycin selection marker. Then, EF1a-dCas9-PRDM9/cdPRDM9-P2A-EGFP-RPBSA-Neo was cloned into a PiggyBac transposon vector (SBI, PB531A-1). sgRNAs were designed using the web-based tool CHOPCHOP. For single sgRNAs, oligos were inserted into pDECKO-mCherry (Addgene, 78534). For multiplex sgRNAs, oligos were first inserted into ph7SK-gRNA (Addgene, 53189), phU6-gRNA (Addgene, 53188), pmU6-gRNA (Addgene, 53187), and phH1-gRNA (Addgene, 53186). Then, four plasmids each carrying single sgRNA were cloned to pLV-GG-hUbC-dsRED (Addgene, 84034) using the NEBridge® Golden Gate Assembly Kit (BsmBI-v2) (NEB).

### Lentivirus generation and transduction

VSV-G, psPAX2, and lentiviral vector containing the sgRNA sequences were co-transfected into 293T cells using Lipofectamine 3000 (Thermo Fisher Scientific) following manufacturer’s instructions. After 48 hours, the culture media was harvested and viral particles were concentrated using PEG-it (SBI). A fluorescence titering assay was performed to measure the multiplicity of infection (MOI) with concentrated virus. An MOI of 10 was used for transduction.

### Generation of transgenic cell lines

Super PiggyBac transposase expression vector (SBI) and PiggyBac transposon vector containing the dCas9-PRDM9 or dCas9-cdPRDM9 cassette were co-transfected into MDA-MB-231 cells by Lipofectamine 3000 (Thermo Fisher Scientific) following the manufacturer’s instructions. After 48 hours, transfected cells were selected with 1 mg/ml G418 for 9 days while passaging every 3 days to establish stable lines. After selection, GFP expression was confirmed by flow cytometry. For co-expression of sgRNAs, lentivirus containing sgRNAs was transduced into dCas9-PRDM9 or dCas9-cdPRDM9 transgenic cells with 8 ug/ml Polybrene (Millipore Sigma). After 72 hours, transduced cells were selected with 1.5 ug/ml Puromycin for 5 days with daily media changes. After selection, mCherry expression was confirmed by flow cytometry.

### Western blotting (WB)

1 × 10^6^ cells were lysed in 100 uL RIPA Buffer (Thermo Fisher Scientific) for 10 min on ice. Samples were centrifuged at 14000 x g for 10 min at 4°C, and soluble protein concentration in the supernatant was measured by Pierce BCA Protein Assay Kit (Thermo Fisher Scientific). 100 ug soluble protein was heated in Laemmli Sample Buffer (Bio-Rad) for 5 min at 95°C. For each well, 20 ug denatured protein was loaded, and the SDS-PAGE gel was run in 1 x Tris/Glycine/SDS Buffer (Bio-Rad) at 200 V for 40 min. Protein in the gel was then transferred to a nitrocellulose membrane in Transfer Buffer (25 mM Tris, 192 mM glycine, 10% methanol, 0.5% SDS) at 250 mA for 1 hour. The membrane was blocked with 5% skim milk for 30 min at room temperature with shaking, and then was incubated with primary antibody overnight at 4°C with shaking. The membrane was washed three times and incubated with secondary antibody for 1 hour at room temperature with shaking. 1 mL SuperSignal West Pico PLUS Chemiluminescent Substrate (Thermo Fisher Scientific) was added to the washed membrane for target protein detection. Chemiluminescence was detected using the FluorChem E (ProteinSimple) documentation system.

### Quantitative PCR (qPCR)

For each reaction well, 10 uL PowerUp SYBR Green PCR Master Mix (Applied Biosystems), 1 uL 10 uM primer mix, 1 uL sample DNA, and 8 uL nuclease-free water were mixed. Three technical replicates were done for each sample in a 96 well plate loaded into QuantStudio 3 Real-Time PCR System (Applied Biosystems). Standard cycling conditions were used: Hold stage (×1): 50°C for 2 min, 95°C for 2min; PCR stage (×40): 95°C for 15 s, 60°C for 1 min. Melt curve stage conditions were: 95°C for 15 s, 60°C for 1 min, 95°C for 15 s. Relative fold change was calculated using the delta–delta Ct method by normalizing to either GAPDH or ACTB.

### ATAC-seq library preparation

5 × 10^4^ cells were harvested for each reaction. ATAC-seq library preparation was carried out using the ATAC-Seq Kit (Active Motif) according to the manufacturer’s instructions.

### Chromatin immunoprecipitation (ChIP)

ChIP was carried out as previously described with some modifications.^33^ For each ChIP reaction, 3 × 10^6^ cells were cross-linked in 1% methanol-free formaldehyde at room temperature for 10 min, and then quenched with 0.114 M Glycine. Fixed cells were resuspended in 300 uL High Salt Lysis/Sonication Buffer (800 mM NaCl, 25 mM Tris pH 7.5, 5 mM EDTA, 1% Triton X-100, 0.1% SDS, 0.5% sodium deoxycholate, 1 x protease inhibitor cocktail) and frozen in -80°C. 20 uL Dynabeads Protein G (Thermo Fisher Scientific) was first incubated with 2 ug antibodies in 20 uL Chromatin Dilution Buffer (25 mM Tris pH 7.5, 5 mM EDTA, 1% Triton X-100, 0.1% SDS, 1 x protease inhibitor cocktail) at room temperature for 3 hours with rotation. Fixed cells were transferred to a Bioruptor Pico Microtube (Diagenode) and sonicated for 10 cycles (30 s on/off) in Bioruptor (Diagenode) at 4°C. 1 mL Chromatin Dilution Buffer was added to sonicated cells, and samples were spun at 13600 x g for 30 min at 4°C to pellet insoluble material. If appropriate, 2 uL SNAP-ChIP spike-ins (EpiCypher) was added to the supernatant. 65 uL soluble chromatin was set aside as input. Dynabead/antibody mix was added to the soluble chromatin and incubated at 4°C overnight with rotation. Then, beads were washed with 1 mL Wash Buffer A (140 mM NaCl, 50 mM HEPES pH 7.9, 1 mM EDTA, 1% Triton X-100, 0.1% SDS, 0.1% sodium deoxycholate), Wash Buffer B (500 mM NaCl, 50 mM HEPES pH 7.9, 1 mM EDTA, 1% Triton X-100, 0.1% SDS, 0.1% sodium deoxycholate), and Wash Buffer C (20 mM Tris pH 7.5, 1 mM EDTA, 250 mM LiCl, 0.5% NP-40 Alternative, 0.5% sodium deoxycholate) once each, and TE Buffer (10 mM Tris pH 7.5, 1 mM EDTA) twice. Bound DNA was eluted twice with 100 uL Elution Buffer (10 mM Tris pH 7.5, 1 mM EDTA, 1% SDS) at 65°C for 5 min. Eluted DNA and input samples were incubated with 4 ug RNase A (Millipore Sigma) at 65°C overnight to reverse cross-linking and digest any contaminating RNA. ChIP and input samples were further digested with 40 ug Proteinase K (NEB) at 45°C for 2 hours and then purified with the Zymo ChIP DNA Clean & Concentrator Kit (Zymo Research) following the manufacturer’s instructions. Purified ChIP and input DNA were used for sequencing library preparation or qPCR. ChIP-seq libraries were prepared using either the KAPA HyperPrep Kit (Roche, KK8504) or the Watchmaker DNA Library Prep Kit (#7K0103-096).

### RNA extraction and reverse transcription

1 × 10^6^ cells were harvested in 1 mL TRIzol reagent (Thermo Fisher Scientific) and homogenized by pipetting. After incubation at room temperature for 5 min, 100 uL BCP was added. Samples were vortexed and incubated briefly at room temperature and centrifuged at 12,000 × g, 15 min, 4°C. The aqueous phase was transferred to a new tube and total RNA was isolated with the RNeasy Micro Plus Kit (Qiagen) following the manufacturer’s instructions. Purified total RNA was used for sequencing library prep or RT-qPCR. For reverse transcription, SuperScript III First-Strand Synthesis Kit (Invitrogen) was used to convert 5 ug purified total RNA to cDNA with Random Hexamer Primer following the manufacturer’s instructions. The resulting cDNA was used for qPCR. Sequencing libraries were prepared from total RNA using the ribosome depletion method with the KAPA RNA HyperPrep Kit with RiboErase (Roche KR1351).

### TT-TimeLapse-seq

TT-TimeLapse-seq was performed as previously described with modifications.^34^ 1 mM 4-thiouridine was added to culture media in a 15-cm dish for 5 min at 37°C. After rinsing twice with ice cold PBS, 1 mL TRIzol reagent (Thermo Fisher Scientific) was added to lyse cells. After incubation at room temperature for 5 min, 200 uL chloroform was added. Samples were vortexed and incubated briefly at room temperature and centrifuged at 12,000 × g, 5 min, 4°C. The aqueous phase was transferred to a new tube and total RNA was isolated by isopropanol precipitation. Genomic DNA was removed using the TURBO DNase Kit (Ambion) and then samples was further cleaned up with RNAclean XP beads (Beckman) following the manufacturer’s instructions. 75 ug purified RNA was sheared in 160 uL RNA fragmentation buffer (75 mM Tris pH 7.4, 112.5 mM KCl, 4.5 mM MgCl2) for 3.5 min at 94°C. Sheared RNA was purified with the RNeasy Mini Kit (Qiagen). For each reaction, 50 ug purified RNA was biotinylated by 5 ug MTSEA-biotin-XX (Biotium) for 30 min at room temperature with rotation. Samples were purified again with RNeasy Mini Kit (Qiagen) to remove unreacted MTSEA-biotin-XX. Purified biotinylated RNA was incubated with 10 uL blocked Dynabeads MyOne Streptavidin C1 beads (Thermo Fisher Scientific) for 15 min at room temperature with rotation. Beads were washed with 100 uL High-salt Wash Buffer (100 mM Tris pH 7.4, 10 mM EDTA, 1M NaCl, 0.05% Tween-20) three times and 100 uL TE Buffer (10 mM Tris pH 7.4, 1 mM EDTA) twice. Nascent transcript was eluted in 25 uL Elution Buffer (100 mM DTT, 20 mM HEPES pH 7.4, 1mM EDTA, 100 mM NaCl, 0.05% Tween-20) for 15 min at room temperature with rotation. 20 uL Eluted RNA was added with 0.835 uL 3M sodium acetate pH 5.2, 0.2 uL 500 mM DETA, 1.3 uL TFEA, 1.365 uL waster, 1.5 uL 200mM mCPBA for TimeLapse chemistry, and then purified with 25 uL RNAclean XP beads (Beckman). 18 uL Purified RNA was added with 0.8 uL water, 0.2 uL 1 M DTT, 0.2 uL 1M Tris pH 7.4, 0.4 uL 0.5 M EDTA, 0.4 uL 5 M NaCl for reducing treatment, and then purified with 20 uL RNAclean XP beads (Beckman). Finally, purified RNA was used for sequencing library prep with the KAPA RNA HyperPrep Kit with RiboErase (Roche KR1351).

### Chromosome conformation capture (3C-qPCR)

3C was done as previously described with some modifications.^35^ For each reaction, 4 × 10^6^ cells were cross-linked in 2% formaldehyde for 10 min at room temperature, and then quenched with 0.135 M Glycine. Cross-linked cells were lysed in 4 mL cold Lysis Buffer (10 mM Tris pH 7.5, 10 mM NaCl, 0.2% NP-40, 1 x protease inhibitor cocktail) for 10 min on ice. Samples were centrifuged for 5 min at 400 x g, 4°C to pellet nuclei. Nuclei were resuspended in 500 uL 1.275x Csp6I restriction enzyme buffer, 7.5 uL 20% SDS was added, and samples were incubated for 1 hour at 37°C with shaking. 50 uL 20% Triton X-100 was added, and samples were incubated for 1 hour at 37°C with shaking. 10 uL aliquot of each sample was taken as undigested genomic DNA control. 400U Csp6I enzyme (Thermo Fisher Scientific) was added, and the sample was incubated overnight at 37°C with shaking. A 10 uL aliquot of each sample was taken as a digested genomic DNA control. 40 uL 20% SDS was added, and the samples were incubated for 30 min at 65°C with shaking. 6.125 mL 1.25 x Ligation Buffer (40 mM Tris pH 7.5, 10 mM DTT, 10 mM MgCl2, 0.5 mM ATP) and 750 uL 10% Triton X-100 were added, and samples were incubated for 1 hour at 37°C with shaking. 20U T4 DNA Ligase (Thermo Fisher Scientific) was added, and samples were incubated for 4 hours at 16°C and for 30 min at room temperature. 300 ug proteinase K was added, and samples were incubated overnight at 65°C. 300 ug RNase A was added, and samples were incubated for 30 min at 37°C. The 3C product was purified by ethanol precipitation and resuspended in 190 uL HindIII restriction enzyme buffer. 100U HindIII (Thermo Fisher Scientific) was added, and the sample was incubated for 2 hours at 37°C with shaking. Digested 3C product and undigested/digested genomic DNA control were purified by ethanol precipitation. The purified undigested/digested genomic DNA control was used for checking digestion efficiency at regions of interest by qPCR. The purified 3C product was used for measuring interaction frequency at target enhancer-promoter pairs by qPCR.

### ATAC-seq and ChIP-seq data analysis

Data were filtered for high-quality reads using the fastq_quality_filter tool from FASTX-Toolkit with parameters -p 80 -q 20 parameters. Filtered libraries were aligned to the hg19 genome assembly using Bowtie2 with parameters --end-to-end --sensitive. Aligned reads were filtered to get uniquely mapping reads without PCR duplicates using SAMtools with parameters -q 1 -f 3 –F 256. Peaks were called using MACS2 with default parameters. Peak intersections were evaluated using BEDTools. Peaks were annotated using the annotatePeaks tool from HOMER. ChIP-seq data across different samples were normalized using CHIPIN with parameters raw_read_count type_norm=“linear”.^36^ Bigwig files were generated using the bamCoverage tool from deepTools. Metagene plots and heatmaps were generated using the computeMatrix and plotHeatmap tools from deepTools with parameters reference-point -b 2000 -a 2000. Correlation plots were generated using the multiBigwigSummary and plotCorrelation tools from deepTools with default parameters.

### RNA-seq data analysis

Data were aligned to the hg19 genome assembly using STAR with default parameters. Aligned reads were counted using HTSeq. Differential gene expression was called using DESeq2 with a cutoff of |Log2_Fold_Change| >= 1.5 and adjusted p-value < 0.01. MA plots were generated using ggplot2.

### TT-TimeLapse-seq data analysis

Data were processed according to the published protocol.^34^ Reads were filtered for unique sequences using FastUniq, trimmed using cutadapt and aligned to the hg19 genome along with annotations using HISAT. Aligned reads were filtered to get uniquely mapping reads using SAMtools. Reads were counted using HTSeq-count using the union mode. To determine the number of uridine residues inferred from each read and the sites of T-to-C mutations, the aligned bam files were processed in R using Rsamtools. Genome tracks were generated using STAR aligner and normalized using factors derived from DESeq2 estimateSizeFactors. Tracks were converted to binary format using the toTDF tool from IGVtools.

## Data availability

Data generated in this study is available through NBCI’s Gene Expression Omnibus under GEO accession number GSE290064. This study also used data from existing ChIP-seq datasets GSE71327, GSE53490, and GSM2572593.

## Acknowledgments

We thank Galina Petukhova for the *Prdm9* cDNA, Nadya Dimitrova for guidance on the 3C-qPCR protocol, Zachary Smith and Yannick Jacob for project advice, Andreas Pintado-Urbanc for guidance on TT-Timelapse-seq, Matthew Simon for TT-Timelapse-seq advice and critical reading of the manuscript, and Emily Forrest for critical reading of the manuscript. We are grateful to the Yale Center for Genomic Analysis for library preparation and sequencing.

## Author contributions

Conceptualization: H.Y., B.J.L.; Investigation: H.Y., Y.Z., Z.L., B.W.W.; Validation: H.Y.; Formal analysis: H.Y.; Visualization: H.Y., B.J.L.; Funding acquisition: B.J.L.; Supervision: B.J.L.; Writing – original draft: H.Y., BJ.L.; Writing – review & editing: H.Y., BJ.L.

