## Supplemental Figures for "H3K4me3 instructs transcription at intergenic active regulatory elements"

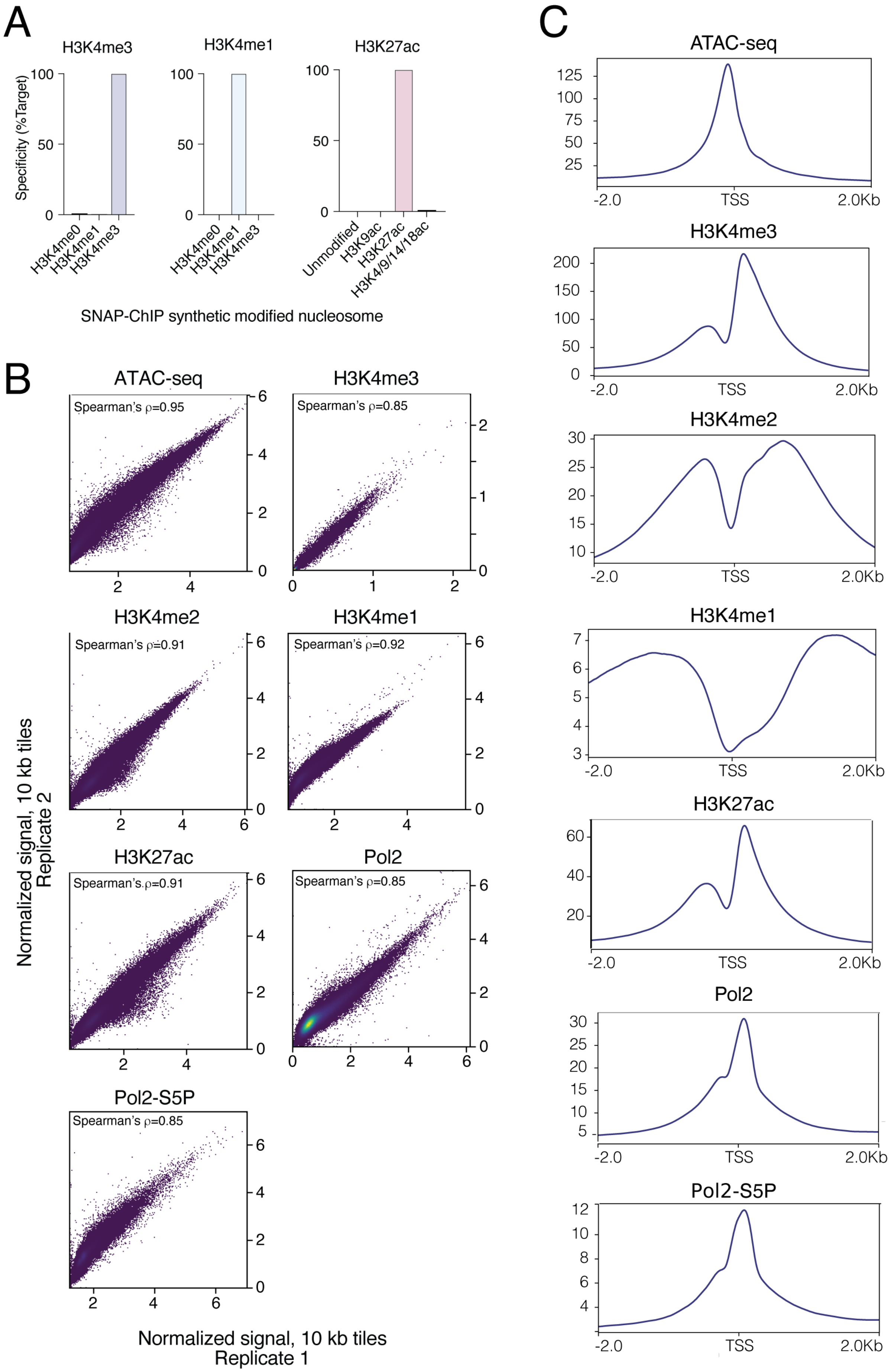


**Supplemental Figure 1. Quality control of ChIP-seq data. A**, On and off target specificities of H3K4me3, H3K4me1, and H3K27ac antibodies measured by ChIP-qPCR with SNAP-ChIP synthetic modified nucleosome spike-ins (Epicypher). **B**, Scatter plots of signal normalized to library size in 10 kb tiles showing correlations between two replicates of ATAC-seq, H3K4me1, H3K4me2, H3K4me3, H3K27ac, total Pol II, and Pol II-S5P ChIP-seq in MDA-MB-231 cells. **C**, Metagene plots of ATAC-seq, H3K4me3, H3K4me1, H3K4me2, H3K27ac, total Pol II, and Pol II-S5P ChIP-seq reads at all promoter transcription start sites (TSSs) confirming expected distributions for each modification.

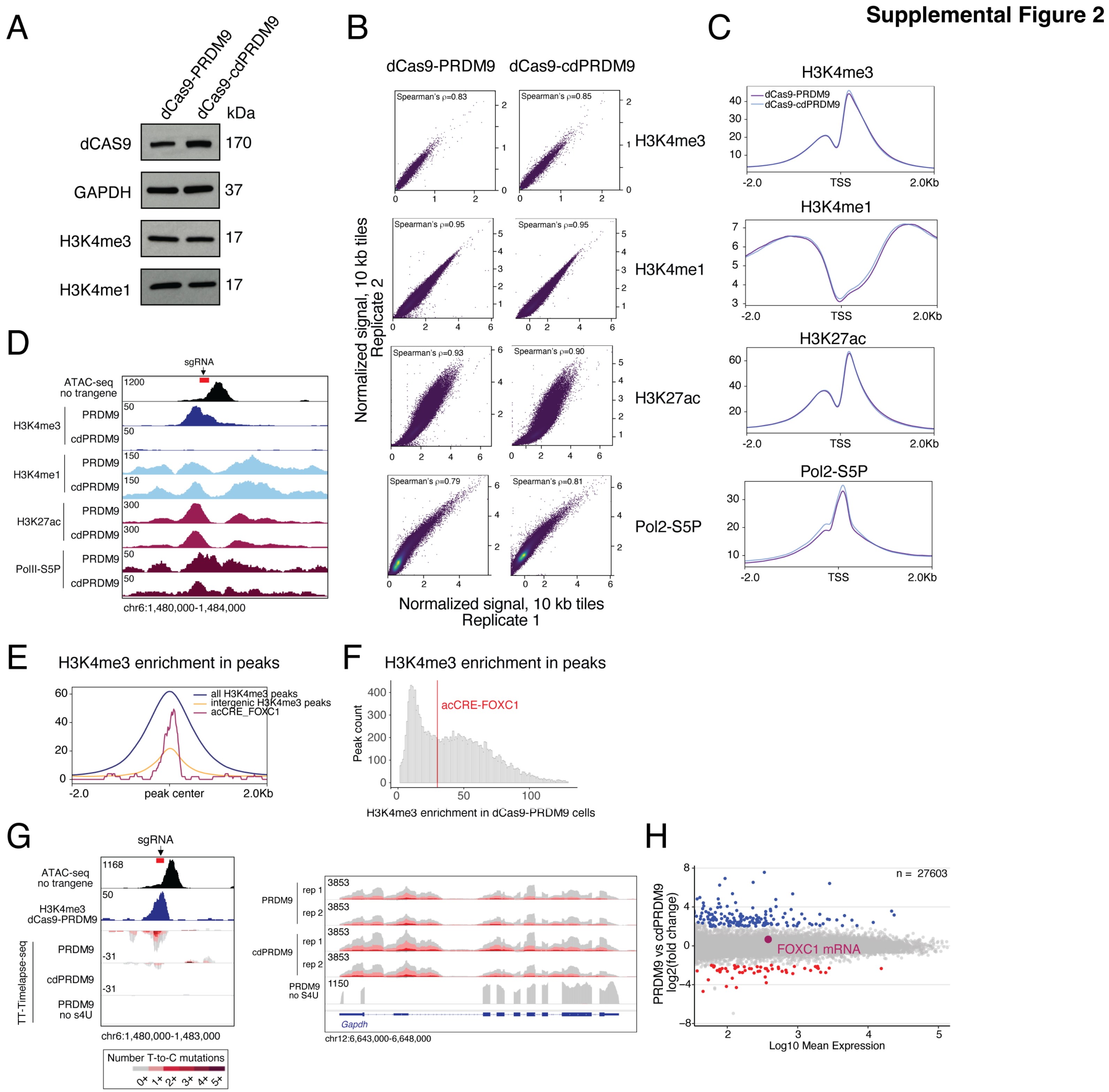


**Supplemental Figure 2. H3K4me3 facilitates local transcription at acCRE_FOXC1. A**, Western blotting for dCas9 (anti-CAS9 antibody), H3K4me3, H3K4me1, and GAPDH in dCas9-PRDM9 and dCas9-cdPRDM9 cells. **B**, Scatter plots of signals in 10 kb tiles showing correlations between two replicates each of H3K4me1, H3K4me3, H3K27ac, and Pol II-S5P ChIP-seq in dCas9-PRDM9 and dCas9-cdPRDM9 cells. **C**, Metagene plots of normalized H3K4me3, H3K4me1, H3K27ac, and Pol II-S5P ChIP-seq reads in dCas9-PRDM9 and dCas9-cdPRDM9 cells at all promoter TSSs. **D**, Genome browser tracks for ATAC-seq in wild type cells, and normalized H3K4me3, H3K4me1, H3K27ac, and Pol II-S5P ChIP-seq signals for the second replicates in dCas9-PRDM9 and dCas9-cdPRDM9 cells at acCRE_FOXC1. Arrow and red bar indicate sgRNA site. First replicates are shown in Figure 2C. **E**, Metagene plot of H3K4me3 ChIP-seq reads in dCas9-PRDM9 cells at all H3K4me3 peaks, intergenic H3K4me3 peaks, and at acCRE_FOXC1 with recruitment of dCas9-PRDM9, showing comparable signal between acCRE_FOXC1 after PRDM9 recruitment and intergenic H3K4me3+ peaks. **F**, Histogram of H3K4me3 ChIP-seq intensity in dCas9-PRDM9 cells among all H3K4me3 peaks. Red line shows the peak intensity at acCRE_FOXC1. **G**, Left, genome browser tracks for ATAC-seq in wild type cells, normalized H3K4me3 ChIP-seq signal, and nascent transcript reads by TT-Timelapse-seq for the second replicates in dCas9-PRDM9 and dCas9-cdPRDM9 cells at the acCRE-FOXC1 locus. Arrow and red bar indicate sgRNA site. First replicates are shown in Figure 2F. Color of the nascent transcript signal stands indicates the number of T-to-C mutations found per read. Right, nascent transcript reads as measured by TT-Timelapse-seq in dCas9-PRDM9 and dCas9-cdPRDM9 cells at the *GAPDH* gene. **H**, MA plot of total nascent transcriptome (TT-Timelapse-seq) reads in dCas9-PRDM9 compared to dCas9-cdPRDM9 (n=2 biological replicates).

**Supplemental Figure 3**

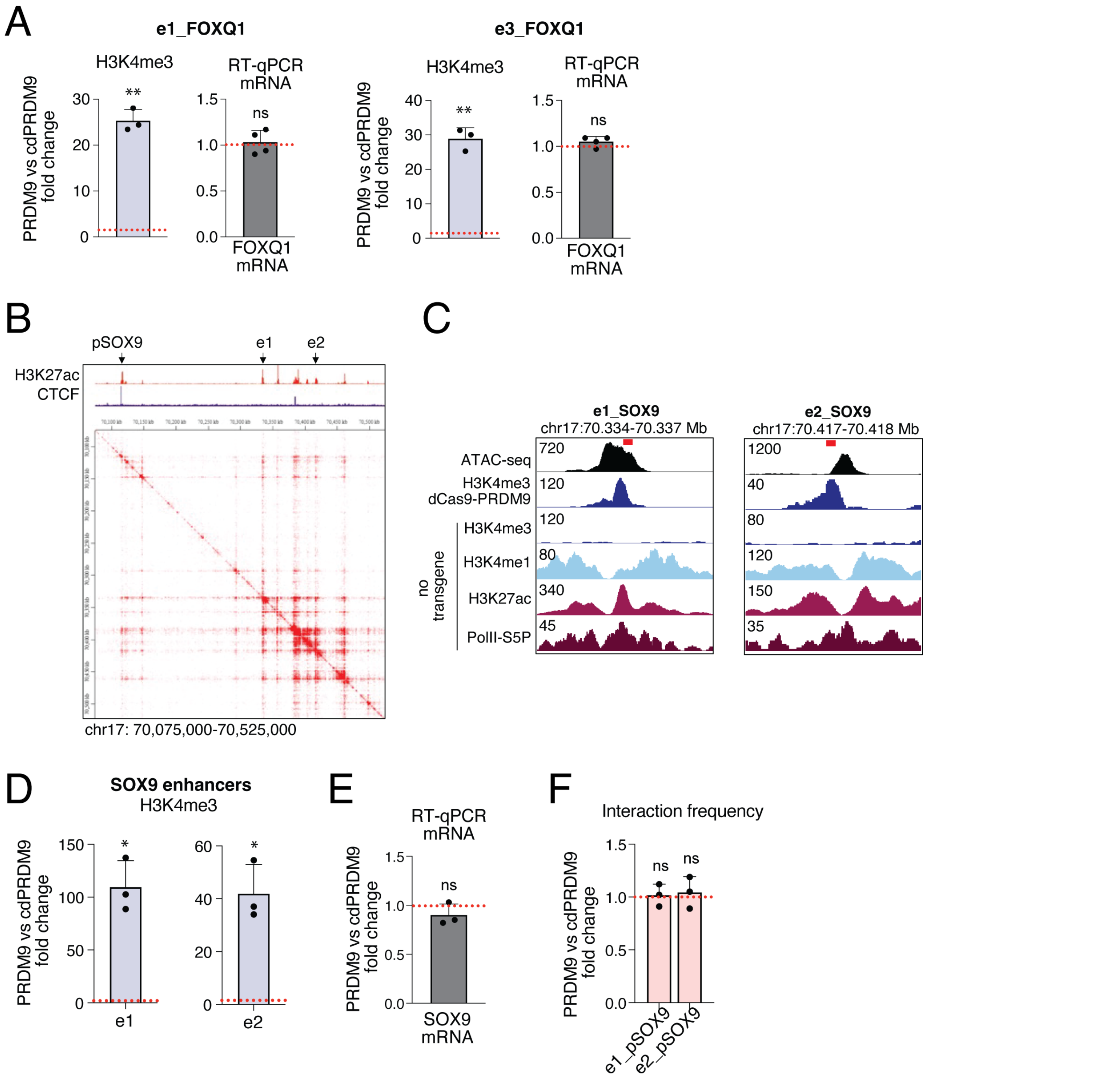


**Supplemental Figure 3. H3K4me3 is dispensable for enhancer activity. A,** Fold change for H3K4me3 levels and FOXQ1 mRNA levels in dCas9-PRDM9 compared to dCas9-cdPRDM9 cells at singly-targeted e1_FOXQ1 and e3_FOXQ1 loci. **P < 0.01, ns, not significant at a threshold of P < 0.05, one-sample t test. **B**, Genome browser tracks of H3K27ac and CTCF ChIP-seq signals (top), and heatmap representing long-range chromatin contact probabilities measured by H3K27ac HiChIP in wild type MDA-MB-231 cells in the region of eSOX9. Arrows point to two target *SOX9* enhancers (e1 and e2) and the *SOX9* gene promoter (pSOX9). Data from GEO:GSM2572593. **C**, Genome browser tracks for ATAC-seq, H3K4me3, H3K4me1, H3K27ac, and Pol II-S5P ChIP-seq signals in wild type cells, and H3K4me3 ChIP-seq signal in dCas9-PRDM9 cells at two target *SOX9* enhancers with expression of multiplexed sgRNAs. Red bars indicate sgRNA sites. **D**, Fold changes of H3K4me3 at two target *SOX9* enhancers in dCas9-PRDM9 compared to dCas9-cdPRDM9 measured by ChIP-qPCR. *P < 0.05, one-sample t test. **E**, Fold change of *SOX9* mRNA levels measured by RT-qPCR. ns, not significant at a threshold of P < 0.05, one-sample t test. **F**, Fold change of normalized interaction frequencies between target enhancers and the *SOX9* promoter measured by 3C-qPCR. ns, not significant at a threshold of P < 0.05, one-sample t test. Error bars represent mean ±SD (n = 3-4 biological replicates).

**Supplemental Figure 4**

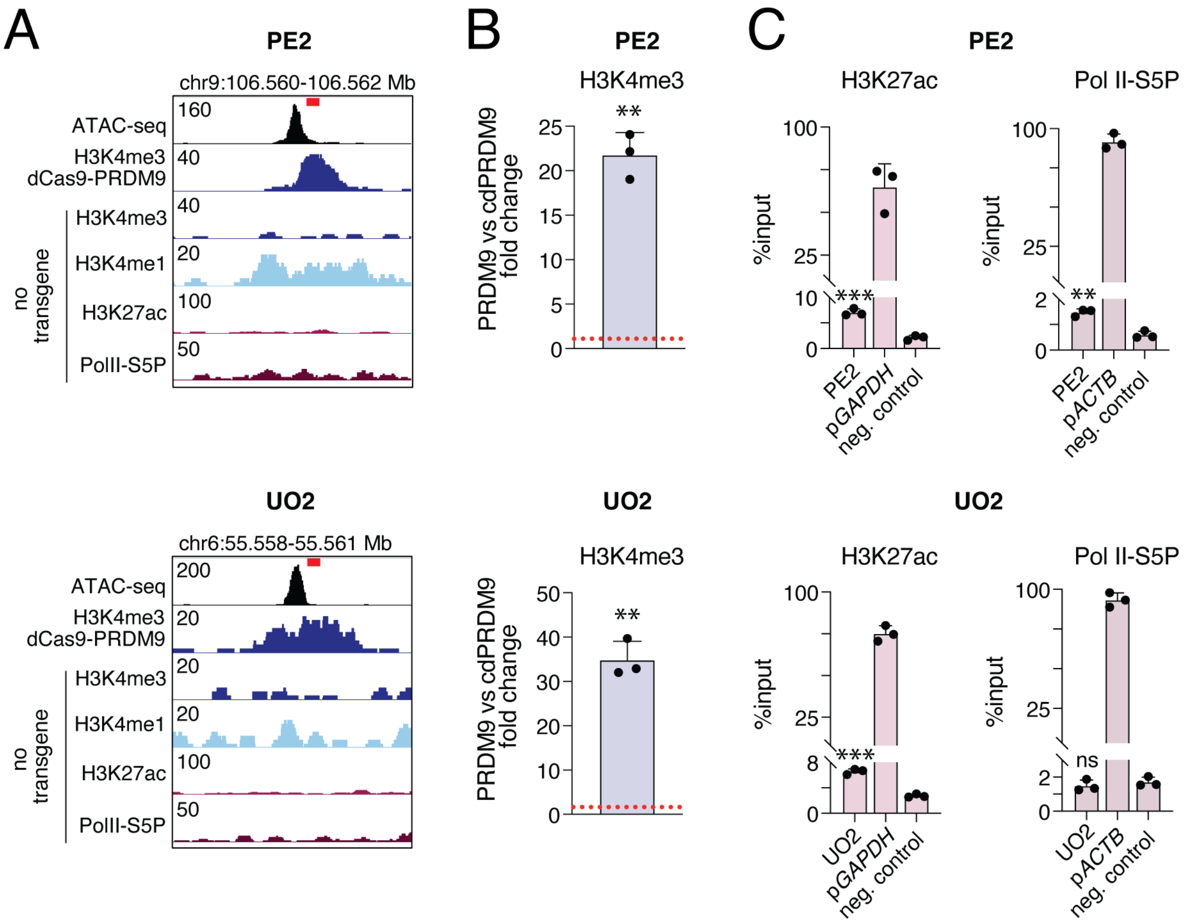


**Supplemental Figure 4. Chromatin and transcriptional changes at an additional primed enhancer (PE) and unmodified open (UO) region. A**, Genome browser tracks for ATAC-seq, H3K4me3, H3K4me1, H3K27ac, and Pol II-S5P ChIP-seq signals in wild type cells, and H3K4me3 ChIP-seq signals in dCas9-PRDM9 cells at the PE2 and UO2 loci. Red bars indicate sgRNA sites. **B**, Fold change in H3K4me3 at PE2 and UO2 loci in dCas9-PRDM9 compared to dCas9-cdPRDM9 cells measured by ChIP-qPCR. **P < 0.01, one-sample t test. **C**, Relative enrichment of H3K27ac and Pol II-S5P at a positive control region (*GAPDH* or *ACTB* gene promoter), a negative control region (gene desert locus on chromosome 12), PE2, and UO2 measured by ChIP-qPCR. ***P < 0.0001, **P < 0.01, ns, not significant at a threshold of P < 0.05, unpaired t-test compared to negative control. Error bars represent mean ±SD (n=3 biological replicates).
